# Acoustic censusing and individual identification of birds in the wild

**DOI:** 10.1101/2021.10.29.466450

**Authors:** Carol L. Bedoya, Laura E. Molles

## Abstract

Avian vocal individuality carries information that can be utilized as an alternative to physical tagging of individuals. However, it is rarely used in conservation tasks despite rapidly-expanding use of passive acoustic monitoring techniques. Reliable acoustic individual recognizers and accurate quantifiers of population size remain elusive, which discourages the use of vocal individuality for monitoring, wildlife management, and ecological research. We propose a neuro-fuzzy framework that allows discrimination of individuals by their calls, the discovery of unexpected individuals in a set of recordings, and estimation of population size using solely sound. Our method, tested using data collected in the wild, allows rapid individual identification and even acoustic censusing without prior information from the recorded individuals. We achieve this by integrating a fuzzy classification and clustering methodology (LAMDA) into a Convolutional Deep Clustering Neural Network (CDCN). Our approach will benefit monitoring for conservation, and paves the way towards robust individual acoustic identification in species whose handling is time-consuming, culturally or ethically problematic, or logistically difficult.

---

Population monitoring is one of the pillars of wildlife conservation (1). It helps determine the processes driving changes in animal communities and populations (2), allows quantification of the effects of disturbance (3), and guides the development of policies and management strategies for habitat and species conservation (1, 4). Nonetheless, wildlife monitoring is a challenging task; some species are cryptic, sensitive to disturbance, difficult to capture and handle, or severely threatened, placing constraints on active monitoring approaches (5).

Vocal individuality, i.e., the set of acoustic features unique to an individual, can be used as an alternative, minimally invasive identification technique (5), and has significant advantages over other commonly-used monitoring approaches (e.g., in situ observation, capture-mark-recapture). Identifying individuals by their calls eliminates the need for making physical contact with the individual (or even having direct line of sight with it), allows efficient long-term monitoring, and reduces personnel costs. However, acoustic individual identification is rarely used in conservation tasks (5, 6).

In comparison to acoustic *species* identification, methods used for *individual* identification require a more refined set of acoustic features to characterize the intra-specific variability of the vocalizations. Since these features tend to be species-specific, individual recognizers are usually tailored for a single species (6), limiting their generalization and application to other taxa. Additionally, individual identification algorithms are usually trained to recognize a pre-defined set of individuals(6–9), and do not allow identification of individuals that were not present during the training stage of the algorithm. This imposes a severe restriction on animal studies in the wild, as the population size (i.e., number of individuals) is usually unknown, and unseen individuals can move and vocalize across wide areas (10, 11), generating unexpected data.

We propose a neuro-fuzzy framework that overcomes previous limitations. Our approach allows identification of avian individuals by their calls, and the discovery of novel individuals that were not present in the training data (i.e., unseen class discovery). Furthermore, it allows determination of the number of individuals in a set of recordings using solely sound (acoustic censusing), which makes it ideal for the monitoring of wildlife populations. Our approach integrates a fuzzy clustering and classification methodology (LAMDA)(12, 13) with Convolutional Neural Networks (CNNs) (14) into a deep clustering framework with the goal of generating a feature set that characterizes vocal individuality (Fig. 1). Then, this feature set is used by LAMDA for acoustic identification and censusing. LAMDA allows supervised, online, and unsupervised learning, and does not require the number of classes as an input parameter (15, 16) when used as clustering algorithm, which provides our framework with capabilities for both unseen individual discovery and acoustic censusing. Moreover, the integration of CNNs in our framework as feature extractors reduces the need for obtaining species-specific acoustic features, making our approach scalable and generalizable to a wide range of species.

**Fig. 1.**
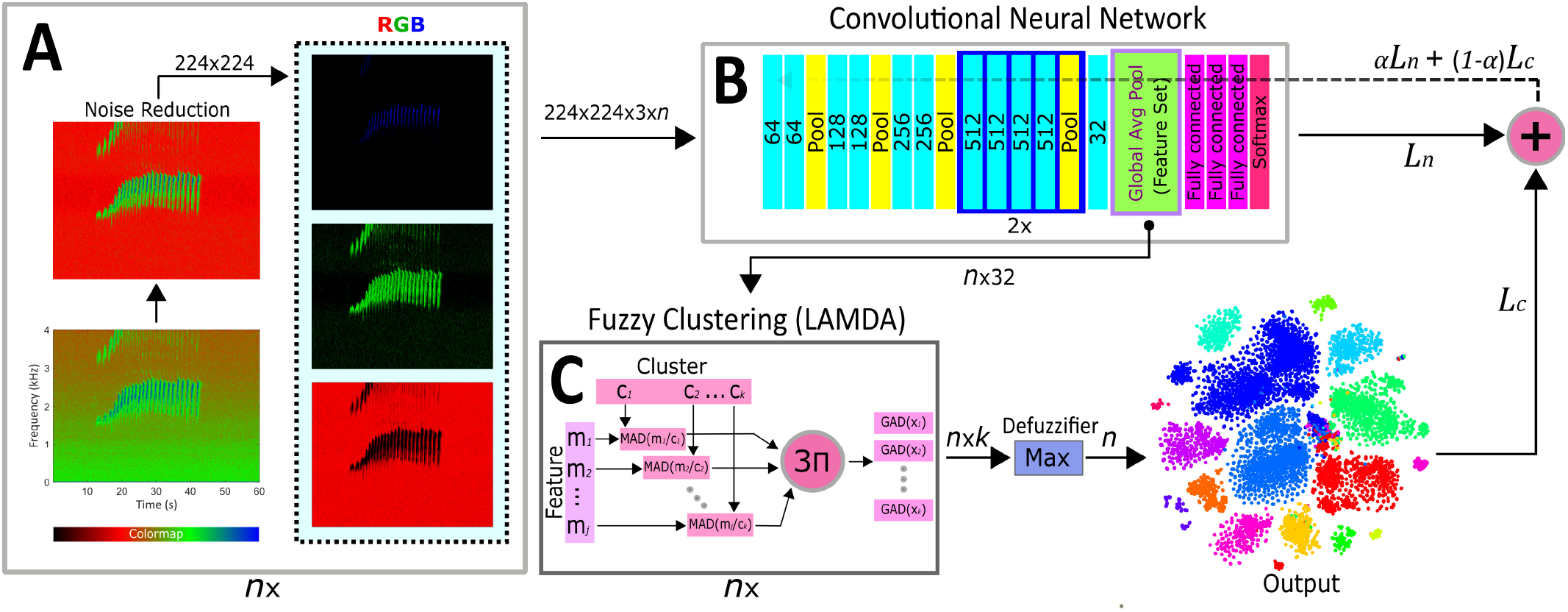
Scheme of the Convolutional Deep Clustering Network (CDCN) used for acoustic individual identification, discovery of unseen individuals, and censusing. The CDCN embeds each call into a feature vector that characterizes vocal individuality, which is then used by LAMDA to identify individuals (known and unexpected) and perform acoustic censusing. The framework uses as input images of the spectrogram of the audio signal, and optimizes both the clustering and classification losses in a joint process. Once the training process has been completed, the training loops are disconnected and the convolutional neural network is used as a feature extractor for LAMDA which performs supervised (classification) and unsupervised (clustering) tasks. Here, *n* is the number of calls to be analyzed, *k* is the number of individuals, *L_n_* and *L_c_* are the network and clustering losses, respectively, and *α* is a mixing factor to combine both losses. Cyan blocks indicate convolutional layers.

We use roroa (also known as great spotted kiwi) *Apteryx maxima* Sclater and Hochstetter, as a model organism to demonstrate the capabilities of our method. Roroa is a nocturnal species that roosts and nests in burrows and its handling presents ethical and logistical challenges; development of a sensitive, minimally-invasive monitoring method will substantially assist their management. In bioacoustics terms, roroa is also an ideal species for testing our method. Roroa individuals typically form highly territorial long-term monogamous pairs, which facilitates tracking and identification, and their large territories (c. 40 ha) reduce the likelihood of neighbouring individuals calling in close proximity to their nests (17, 18). Both sexes produce stable and loud well-defined calls with inter- and intra-sexual vocal variability (19). These calls are comprised of series of similarly-shaped syllables (20), which facilitates automated analysis and comparison.

We propose a paradigm shift in the analysis of bioacoustic data that significantly facilitates the identification and monitoring of populations and individuals in the wild. Our approach addresses a grand challenge in ecoacoustics – the acoustic censusing of animal populations – while significantly contributing to the broader areas of ecology and animal conservation.

## A Convolutional Neural Network for Acoustic Individual Identification

A CNN (Fig. 1B) was designed to analyze acoustic data and identify individuals by their calls. The network uses as a core the architecture VGG19 (21), which was modified to improve the regularization of the latent space when used as a feature extractor. One of the advantages of using CNNs in our framework is that they are intrinsically designed to extract features relevant to their task. This removes the need to manually select a set of acoustic features for the species, which is a significant bottleneck in the generalization of individual identification approaches. We chose VGG19 because of its high individual identification accuracy (*SI Appendix, Fig. S1*) and its relatively shallow architecture, as very deep CNNs have regularization issues when integrated into our deep clustering framework. Acoustic data were pre-processed (segmented, augmented, and noise-reduced, see Fig. 1A and methods) before subsequent analysis by the CNN as RGB images. Our network, which we call KiwiNet (Fig. 1B, see methods and *SI Appendix, Table S1*, for an extended description), attained better performance metrics than most of the commonly-used CNN architectures (*SI Appendix, Fig. S1*), with the benefit of having additional layers that help regularize the latent space for clustering analyses.

The differences between KiwiNet and VGG19 are (i) a convolutional layer before the fully connected layers to reduce the number of filters (from 512 to 32) and (ii) a global average pooling layer to embed the call characteristics into a 32-element feature set (latent space), which is also the input feature vector for the clustering stage (Fig. 1C). Imposing a strong bottleneck during feature extraction helps the performance of the clustering algorithm, as analyzing data in a high number of dimensions tends to deteriorate the clustering accuracy (22, 23) and ultimately hinders the regularization of the CDCN during the joint training (clustering + CNN, Fig. 1).

Data is analyzed by the CNN using a colormap that correlates the image colors with the levels of intensity in the spectrogram (Fig. 1A, *SI Code*). We chose KRGB as it avoids color mixtures that could disrupt the spectrogram’s power representation. It also allows specific sections of the call to be independently processed by different convolutional kernels as they are located in separate image channels. In our colormap, background noise is concentrated predominantly in the red channel while the individual’s acoustic information is mostly in the green and blue channels (Fig. 1A). During the pre-processing stage, we augmented the data with background sounds from several sites and recorders in order to minimize confounding factors from the environment and recording equipment in the individual identification process. We also applied a median equalizer after spectrogram estimation (Fig. 1A, see methods) to noise-reduce the data and increase the identification accuracy. The backbone of KiwiNet (VGG19) was pre-trained with the imagenet dataset (24) to both accelerate the training workflow and improve generalization using transfer learning.

## A Neuro-fuzzy Approach for the Optimization of the Acoustic Feature Set

In order to provide our framework with capabilities for recognizing unseen individuals and performing acoustic censusing, we integrated LAMDA, a fuzzy clustering and classification methodology (Fig. 1C), into a convolutional deep clustering network (CDCN). In the CDCN, the latent space of the CNN (Fig. 1B) is used as the input feature set for LAMDA, which analyses the data in *online learning* mode (i.e., cluster prototypes for the known individuals are generated using the data labels, but new clusters can be created). Then, clustering and network losses, *L_c_* and *L_n_*, are linearly combined (*αL_n_* + (1 – *α*)*L_c_*) and back-propagated throughout the CNN (Fig. 1).

The training process of the CDCN (Fig. 1) consists in pretraining KiwiNet to identify the calls of a set of individuals (e.g., 10 ♂). Then, KiwiNet is connected to LAMDA, which uses its latent space as the input feature set. Each iteration, LAMDA assigns each individuals’ calls to their respective clusters using online learning. This assignment is then compared with the data labels using a cross-entropy function, whose result is aggregated to the CNN loss to be jointly back-propagated throughout the CNN via stochastic gradient descent with momentum and restarts. The loss of the CNN is never removed from the CDCN training loop so that LAMDA cannot completely collapse the initial feature embedding.

Jointly training the CNN with a clustering algorithm produces a feature set that accurately characterizes vocal individuality and is *useful for clustering tasks* (e.g., censusing). When used for individual identification, these deep features significantly outperform other commonly-used acoustic descriptors of animal vocalizations (e.g., spectro-temporal features, MFCCs; Fig. 2), as the CDCN optimizes its feature extraction process to generate highly compact clusters. Once the CNN has been jointly trained with the clustering algorithm, the training loops are disconnected and the CNN is used solely as a feature extractor for LAMDA, which performs acoustic individual classification (supervised learning), unseen individual discovery (online learning), and acoustic censusing (unsupervised learning).

**Fig. 2.**
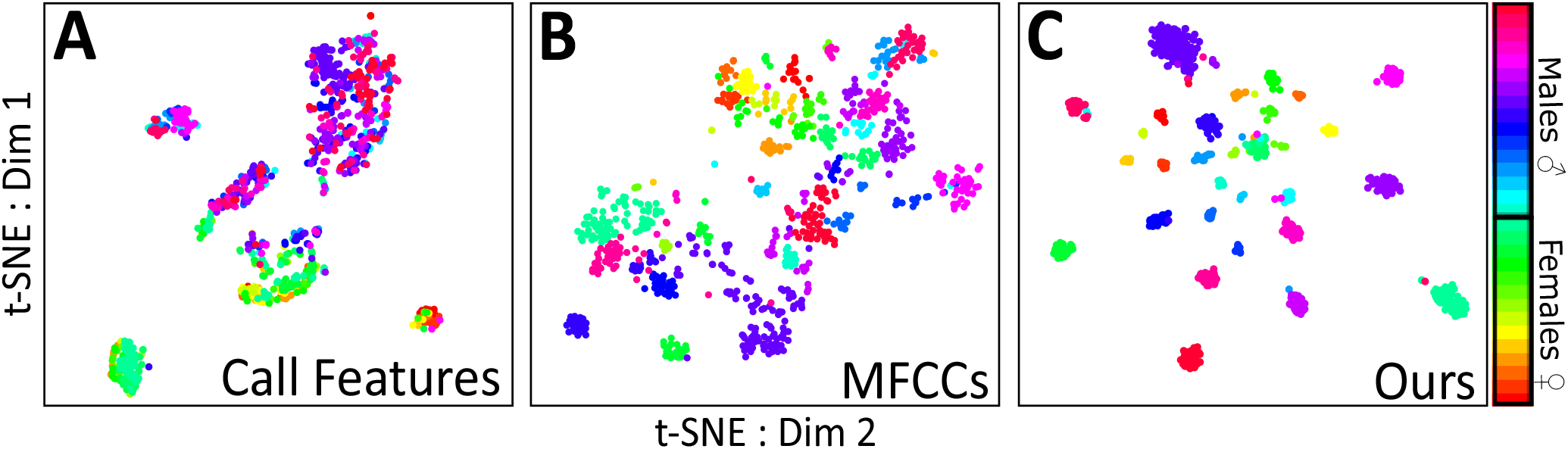
Comparison of our proposed feature extraction method with two common feature sets used in bioacoustics and ecoacoustics. Plots (A-C) are 2D t-SNE ordinations generated using calls from 30 individuals (16 ♂ and 14 ♀). (**A**) Spectro-temporal call features (call duration, syllable duration, inter-syllable interval, number of syllables, dominant frequency, spectral centroid, bandwidth). (**B**) Mel-Frequency Cepstral Coefficients (MFCCs). (**C**) Deep features extracted using our proposed convolutional deep clustering network. Each color represents a different individual.

## Acoustic Individual Identification and Unseen Individual Discovery

After training the CDCN, optimization loops are disconnected and LAMDA can be used in supervised mode to identify known individuals, i.e., cluster prototypes are determined *a priori* from the training dataset and used to classify call features extracted via KiwiNet. On the other hand, to recognize individuals that were not present during the training stage, a common scenario with mobile species or at the borders of individuals’ territories, LAMDA is used in online-learning mode. In this mode, new clusters are created when the highest membership degree of a call belongs to a non-informative class ((13), methods). The prototypes of the original set of clusters (i.e., known individuals) are left intact during the clustering analysis, as these were found supervisedly and are robust (i.e., training accuracy > 90%), and only the prototypes of the recently created clusters are updated as new data is grouped into them. LAMDA, in online-learning and unsupervised modes, tends to generate a number of new clusters that is exact or similar to the number of expected unknown individuals. Nonetheless, a cluster validation stage (methods) was implemented to detect disjoint or sparse clusters that cannot be confidently associated with a specific individual.

In our experiment, we used LAMDA to identify the calls of 20 known (10 ♂ and 10 ♀) and six unknown (3 ♂ and 3 ♀) individuals. Since our model organism (roroa) exhibits significant vocal differences between males and females, we independently trained two CDCNs to recognize calls from each sex. Our method identified calls from known individuals with average accuracies of 89.4%(♂) and 94.6%(♀) (Figs. 3C-F, *SI Appendix, Table S2*), created six high-confidence clusters corresponding to the unknown individuals (Figs. 3A,B), and accurately grouped unknown individuals’ calls into the appropriate clusters (Figs. 3C,D). Our approach not only allows detection and quantification of unseen individuals, but also the matching of each call with its respective singer, which is a never-before-seen way of performing passive monitoring of animals. The identification accuracy of our approach is not as high as that of fully-supervised CNNs (*SI Appendix, Fig. S1*), but allows acoustic monitoring of individuals that have not been marked, or even detected, using other field methods.

**Fig. 3.**
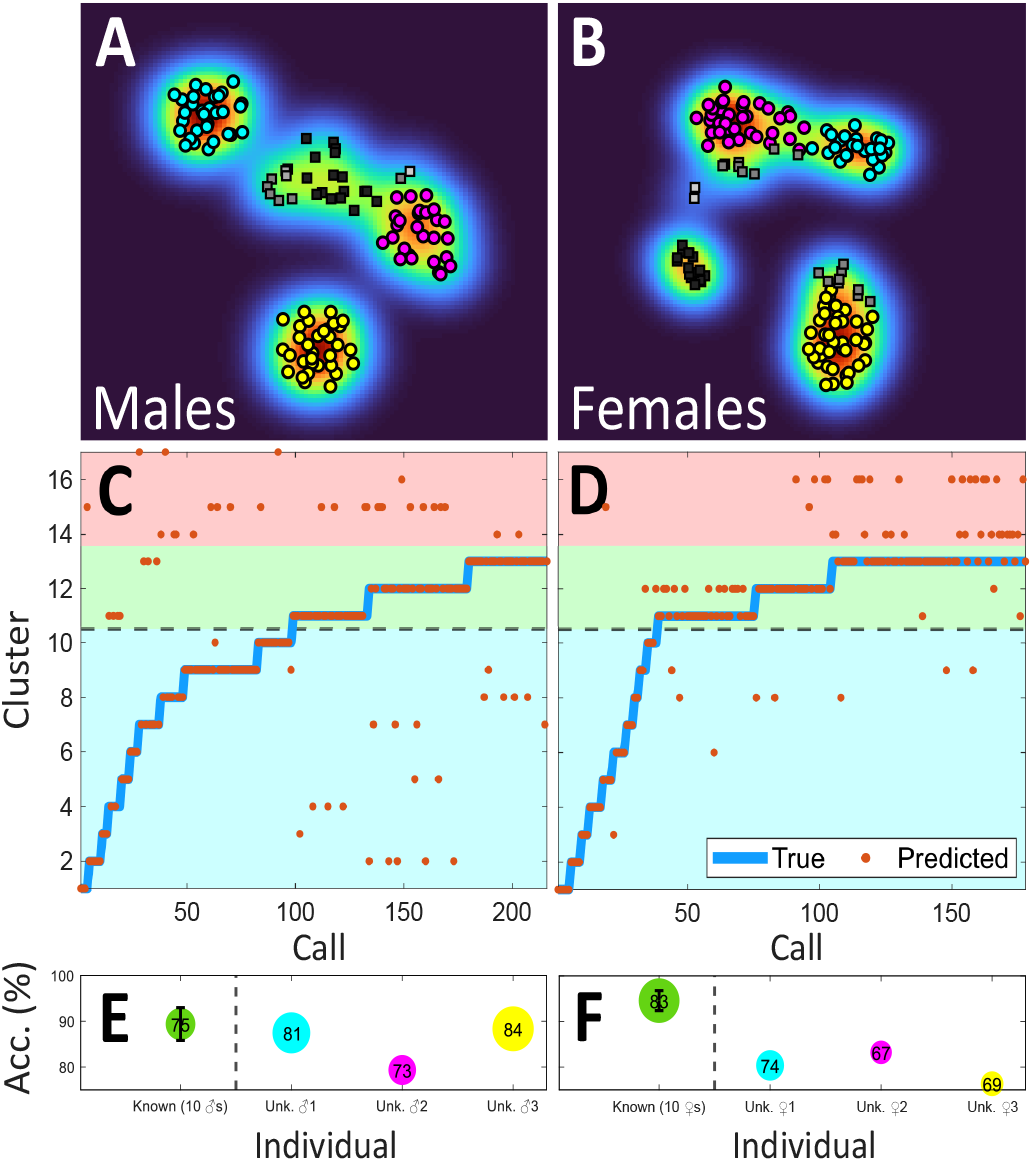
(**A-B**) 2D t-SNE visualization of the new clusters generated by LAMDA in online learning mode. The ordination was performed on the feature set extracted via KiwiNet. Markers in color represent calls from six individuals absent during the training stage of the convolutional deep clustering network (three ♂ (**A**) and three ♀ (**B**)). Data with gray-scale and rectangular markers indicate calls that were categorized as low-confidence clusters (see methods and *SI Appendix, Fig. S2*). The colormap illustrates the data density. t-SNE is used only for visualization and it is not part of the data analysis process. (**C-D**) Identification results for known and unknown males (**C**) and females (**D**). Each orange dot represents a single call (predicted class), which is assigned to one either known or newly-generated cluster. Blue lines indicate the data labels for known and unknown individuals (true class). Clusters in the blue region indicate validation data from known individuals; the green region contains clusters corresponding to individuals unseen during the training stage of the algorithm (clusters in color in **A-B**); and the red region encloses low-confidence clusters that were disregarded during the cluster validation stage (gray-scale clusters in **A-B**). (**E-F**) Identification accuracy (sensitivity/2 + specificity/2) per individual for both males (**E**) and females (**F**). Position of the marker indicates the validation accuracy; size of the marker and the number inside it represents the F1 score (2(precision×sensitivity)/(precision+sensitivity)); color of the marker matches the color of the clusters in Figs. (**A-B**). Dotted lines in (**C-D-E-F**) separate known and unknown individuals. The first marker in both (**E**) and (**F**) is the average accuracy of the known individuals (10 ♂ (**E**) and 10 ♀ (**F**)), where the error bar indicates the standard error. A detailed description of the identification accuracies for each individual (known and unknown) is presented in SI *Appendix (Table S2*).

## Acoustic Censusing

Acoustic censusing - estimating the number of individuals in a set of calls entirely from previously-unseen singers - is performed using LAMDA in unsupervised mode (clustering), where data are grouped based on the fuzzy adequacy degrees of the acoustic features extracted via the KiwiNet trained in the CDCN. LAMDA generates clusters as data are analyzed by the algorithm; consequently, it does not require a pre-defined number of clusters as an input parameter. This approach circumvents the need to have prior knowledge of population size, which has been a barrier to acoustic censusing of animal groups. A second obstacle to censusing via clustering is achievement of a consistent one-to-one correspondence between the number of individuals and the number of generated clusters. We address this by implementing several strategies to reduce cluster over- and under- generation: (i) using CNNs as feature extractors; (ii) performing joint loss optimization with the clustering algorithm and the CNN; (iii) augmenting and noise-reducing the data; (iv) selecting an adequate CNN for acoustic feature embedding (v) using aggregation operators that naturally restrict the number of generated clusters (16); and (vi) implementing a cluster validation stage. Our approach allows accurate individual identification, and it is a true estimator of the number of individuals present in a set of recordings. This significantly differs from most current acoustic-based methodologies for population monitoring, which primarily focus on determining presence/absence and call rates (25, 26), or use indices based on the distribution of energy in the acoustic spectrum (27–29).

A further advantage of addressing acoustic censusing as a clustering problem is that it assigns each call to its respective individual in an unsupervised way, allowing not only quantification but also monitoring of unknown individuals. Figure 4 shows the results of our acoustic censusing methodology applied to 10 unknown roroa individuals (i.e., not used to optimize the feature extractor). We trained two CDCNs on 20 individuals (10 ♂ and 10 ♀, 10 individuals each), to generate two feature extractors that accurately characterize vocal individuality for each sex. Then, we used these features to cluster the calls from 10 new individuals (6 ♂ and 4 ♀). Our method generated 10 high-confidence clusters (Figures 4A-D) corresponding to the 10 expected individuals. Call assignment to these clusters had average accuracies of 88.2%(♂) and 88.6%(♀) (Figs. 4E-F, *SI Appendix, Table S3*), which is remarkable considering this task was performed unsupervisedly with no training data from these individuals.

**Fig. 4.**
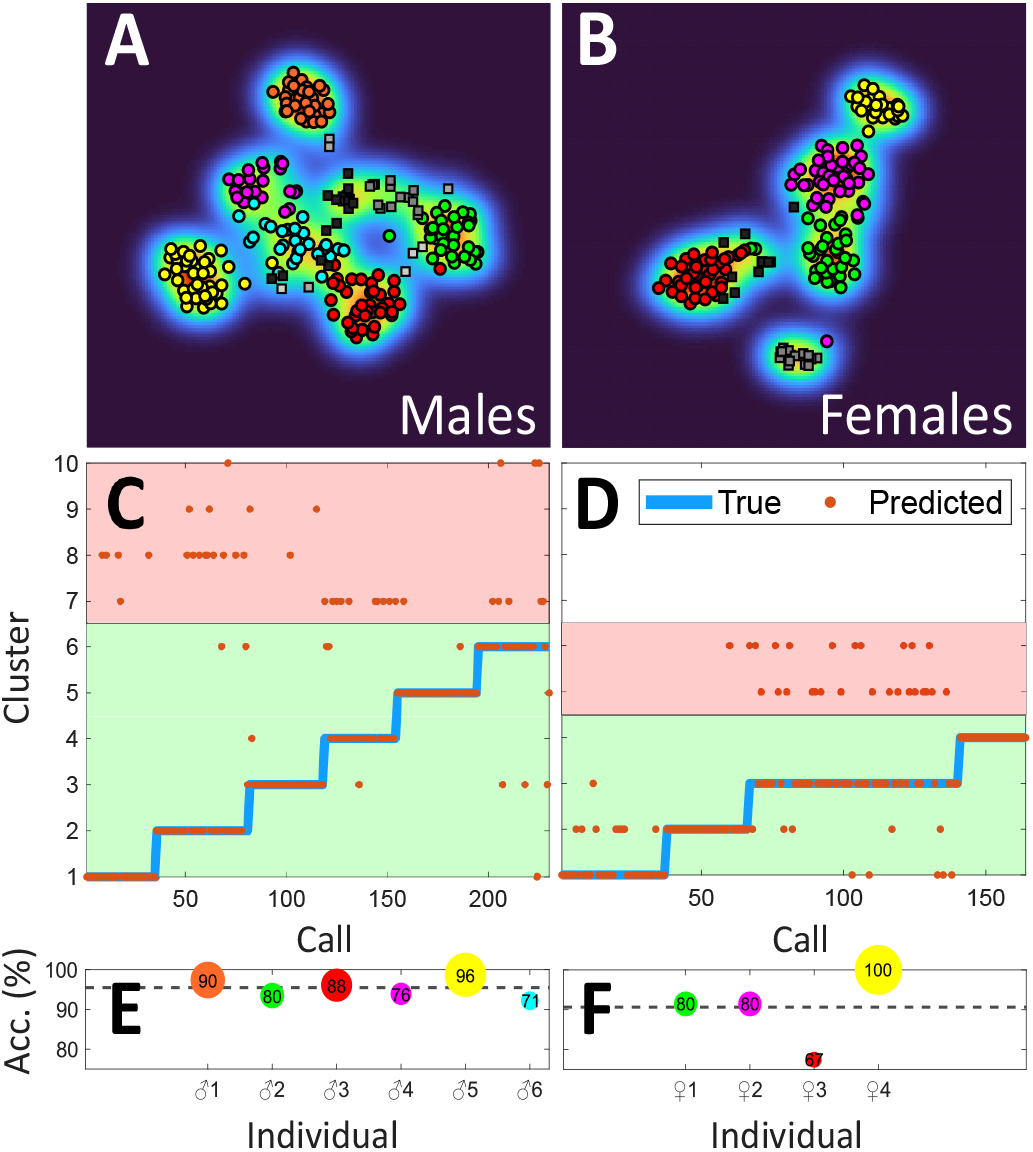
(**A-B**) 2D t-SNE visualization of the calls from 10 individuals (six ♂ **A**, and four ♀ **B**) used to test the acoustic censusing capabilities of our approach. Acoustic censusing is performed by LAMDA in unsupervised mode (clustering) on the features extracted with KiwiNet (after joint training with LAMDA in the CDCN framework). Markers in color indicate clusters that can be confidently associated with an individual. Data with gray-scale and rectangular markers indicate low-confidence clusters rejected during the cluster validation stage (see methods and *SI Appendix, Fig. S2*). The colormap illustrates the data density. t-SNE is used only for visualization and it is not part of the data analysis process. (**C-D**) Censusing results for males (**C**) and females (**D**). Each orange dot represents a call, which LAMDA assigns to a specific cluster (predicted class). Blue lines (true class) are the data labels (these are unseen by the algorithm and only used for visual comparison). The green area encloses the clusters that can be confidently considered individuals, the red area contains the low-confidence clusters disregarded during the cluster validation stage (gray-scale markers in **A-B**). (**E-F**) Accuracy (sensitivity/2 + specificity/2) results for each censused individual male (**E**) and female (**F**). The average censusing accuracy of our approach was 88.2% ± 7.0% for males and 88.6% ± 10.7% for females (mean ± SD, mean indicated by the dotted line in **E-F**). The size of the markers and the numbers inside them indicate the F1 score (2(precision×sensitivity)/(precision+sensitivity)) for each cluster. Colors in Figs. (**E-F**) correspond to the clusters in Figs. (**A-B**). A detailed description of the identification accuracies for each censused individual is presented in SI *Appendix (Table S3*).

## Conclusions

We introduce a fuzzy-based deep learning approach for the classification and clustering of bioacoustic data. The core of our approach is a fuzzy clustering and classification methodology (LAMDA), which is jointly trained with a convolutional neural network to generate a feature set that characterizes the intraspecific variability of vocal individuality. These features are then used by LAMDA to identify individuals by their calls, detect unexpected individuals that were not present in the training stage of the algorithm, and perform acoustic censusing.

This is the first time that LAMDA has been integrated into a deep learning framework, providing embedding capabilities to an already powerful machine learning technique. Our neuro-fuzzy approach opens a new avenue for research in the field of deep clustering, while addressing significant challenges in ecology and animal conservation.

In ecological terms, the most important characteristic of our framework is that it allows quantification of the number of individuals in a set of recordings, which is a rapid and non-invasive approach for the estimation of population size. Nowadays, acoustic indices and call counts are used as proxies for abundance and other biodiversity metrics; however, accurate acoustic methods for the census of individuals remain elusive. Our framework is a significant step in that direction, permitting not only accurate quantification of individuals, but also the unsupervised assignment of calls to their respective sound sources.

Our framework was tested in roroa; however, the use of CNNs in our approach reduces the need to manually curate a specific set of features for the species (e.g., call duration, dominant frequency), and it can be directly applied to sounds from other species with low acoustic complexity. In our case study, calls of both male and female roroa were analyzed using exactly the same parameters despite the sexes having distinctive vocalizations (*SI Data*). Our next step is providing our framework with capabilities for sequential, or syllable-by-syllable, analysis, so we can extend this approach to species with more complex calls and/or call repertoires.

We propose a novel way to census populations acoustically and monitor known and unexpected individuals over time. This will allow ecologists and ethologists to tackle questions related to communication networks, social systems, and behavior in the wild that were previously intractable. Our framework also facilitates monitoring and conservation tasks, especially in species that are cryptic, logistically difficult to capture and tag, or whose handling raises ethical or cultural concerns. Our work challenges current approaches for the analysis of acoustic information in ecoacoustics, ornithology, and animal communication, while simplifying the monitoring and conservation of avian populations in the wild.

## SI Datasets and Algorithms

Acoustic datasets and Algorithms used in this manuscript are available at: https://doi.org/10.6084/m9.figshare.16850542.v1 http://github.com/carolbedoya/Bird-ID-and-Censusing

## Materials and Methods

### Data Collection

Calls of male and female roroa (great spotted kiwi) were collected in the Paparoa mountain range (171.375174, - 42.347427, Blackball – New Zealand), Hawdon Valley (171.743907, −42.961614, Arthur’s Pass National Park – New Zealand), and Nina Valley (172.331246, −42.462351, Lake Sumner Forest Park – New Zealand) during the 2020/2021, 2012-2015, and 2015-2017 breeding seasons, respectively. In total, vocalizations from 30 individuals (8 ♂ and 6 ♀ from the Paparoa Range, and 8 ♂ and 8 ♀ from Arthur’s Pass/Nina Valley) were used to develop and demonstrate the capabilities of our framework. A further 6 individuals were recorded, but excluded from analyses because few calls were captured (< 8 per individual). For 11 of the individuals used in the analysis, calls were collected during two or more breeding seasons; this includes two individuals who were recorded in Nina Valley following their translocation from Hawdon Valley to Nina Valley between the 2014/15 and 2015/16 breeding seasons. A total of 54 autonomous recorders were used at 28 nests across all sites and seasons (AudioMoth v1.1.0 – Conservation Technology Limited, 16 kHz sampling rate and 16-bit depth (Paparoa); AR3 (Hawdon) and AR4 (Nina and Paparoa) – New Zealand Department of Conservation, 8 kHz sampling rate and 16-bit depth; SoundCache (2 site-years in the Hawdon – Cornell Laboratory of Ornithology, 22.05 kHz sampling rate and 16-bit depth). Multiple recorders were placed at some nest sites.

Recorders were placed in the vicinity of roroa nests where at least one member of the pair was fitted with an activity-logging radio transmitter; sharp drops in activity levels for multiple days characterise incubation by males, who typically perform the bulk of incubation, including all daytime incubation (30). Nests were located by using radio-tracking to determine the daytime locations of incubating males. In most cases nests were found using a close-approach technique, which involves spiralling in on a transmitter signal until the location has been encircled. Locations of four nests which could not be closely approached were estimated by triangulating incubating males’ transmitters. Nests within a site-year were on average 3445.9 ± 2144.6 m (mean ± SD) apart, with the minimum distance between any two recorded nests being 419.3 m. Recorders were placed on average 110.8 ± 105.6 m (mean ± SD) away from their target nest sites, with 18 of 28 nests having at least one recorder within 50m. Recorders operated during the night-time period when roroa are expected to be active (30 m after sunset – 30 m before sunrise).

### Call Segmentation

Calls were automatically segmented using a CNN (AlexNet) trained to recognize male calls, female calls, duets and background noise. The procedure consisted of analyzing the spectrogram of 1-min sections of every recording with the CNN in order to detect the presence and sex of an individual. All recordings were downsampled to 8kHz in order to standardize the dataset. Once the presence of a call was detected, the time frame was expanded to three minutes, and analyzed every second by the CNN to refine boundaries of the segmentation. Segments where calls were not detected were also stored for a subsequent use in the data augmentation stage. Data for training the CNN for segmentation were obtained by manually searching a subset of recordings divided into continuous 1-min sections; the resulting training set consisted of 1000 calls from females, 1000 calls from males, and 20000 noise samples, divided in proportions 80/20 for training and validation. For segmentation training and all subsequent analyses, we excluded duets in which male and female roroa overlapped or interspersed their syllables so that each segment could be unequivocally assigned to a single individual.

### Data Pre-processing

Segmented calls were assigned labels based on the nest location associated with the recorder. To minimise the risk of including calls from neighbouring individuals, only calls with signal-to-noise ratios (SNRs) superior to 6 dB (i.e., four times the power of the background noise) were considered for data analyses. In the rare cases in which a call was simultaneously acquired with an SNR > 6 dB by multiple recorders, only the call with the largest SNR was used. The final dataset comprised 849 total calls across the 30 included individuals.

### Acoustic Datasets

The acoustic data from the 30 individuals were divided in four subsets: (i) calls from 20 individuals (10 ♂ and 10 ♀, 319 calls) were used to train both CNN (KiwiNet) and CDCN; (ii) a validation subset (136 calls) with additional calls from those 20 individuals to test KiwiNet’s identification accuracy and the capabilities of our framework to recognise known individuals; (iii) calls from six unseen individuals (3 ♂ and 3 ♀, 258 calls) to evidence the capability of our framework to recognize individuals that were not present during the training stage of the algorithm; and (iv) the remaining four individuals (3 ♂ and 1 ♀, 136 calls) were used in conjunction with the six individuals from (iii) in an entirely unsupervised fashion to test our framework’s ability to perform acoustic censusing. Note that (ii), (iii), and (iv) were run independently, rather than sequentially, on a network pre-trained using solely the original set of training calls (i).

### Data Augmentation

In order to avoid reporting overoptimistic results, the data were augmented by mixing every call with background sounds from different sites. Each call was mixed (constant energy pan: 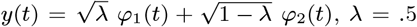) with a random recording (sampling without replacement) from a subset of 1000 background sounds per site (i.e., stratified data augmentation (6)). This was done to reduce the experimental cofounding effects of background sounds. This procedure increases the size of the dataset from *k* to *n* × *k*, where *n* is the number of sites and *k* is the number of recordings. We found that *n* = 6 was a good memory/training-time compromise for our computational setup. The noise segments used for data augmentation are included in *SI Data*.

### Spectrogram Estimation

Spectrograms were computed using a Hann window of 1024 samples with 768 overlap, and 1024 frequency bins. Spectrograms were then noise-reduced and plotted using a KRGB (Black-Red-Green-Blue) colormap (Fig. 1A, *SI Code*) and stored as images before their subsequent analysis by the CNN.

### Noise Reduction

Background noise reduction was performed using a median equalization on the spectrogram of each recording (*SI Code*). This method does not require any tuning parameters, such as cut-off frequencies or thresholds. Let ***X*** = [***x*_1_, *x*_2_, …, *x_f_*, …, *x_N_f__***]^***T***^ be a *N_f_* × *N_t_* matrix representing the magnitude squared of the short term FFT, where 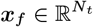 is a vector representing the frequency bin *f*. Thus, the equalization is performed using:

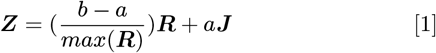

where 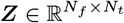 is the noise-reduced version of the magnitude spectrogram, 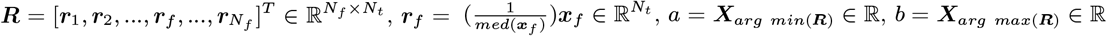 and 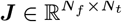 is an all-ones matrix. Magnitude spectrograms (***X***) were softened using a one-dimensional median filter (5 × 1) before the median equalization. After estimating ***Z***, it was converted to decibel units (***Z*_dB_** = 10 · ***log*_10_(*Z*)**) and bilinearly interpolated so that each output pixel value was a weighted average of the pixels in the nearest 2-by-2 neighborhood. This process further noise-reduces the spectrogram and matches the dimensions of ***Z*_dB_** with the input dimensions of the CNN.

### Convolutional Neural Network (CNN)

We used KiwiNet (*SI Appendix, Table S1, SI Code*) as the feature extractor. KiwiNet has a similar architecture to VGG19(21), but introduces before the fully-connected layers: (i) a convolutional layer to reduce the number of filters from 512 to 32 and (ii) a global average pooling layer to generate a 1-dimensional latent space. Two CNNs were pre-trained (one for each sex) using data from roroa individuals (10 ♂ and 10 ♀) before the joint training with the clustering algorithm in the CDCN. The pre-training was performed using stochastic gradient descent with momentum in mini-batches of eight calls with a learning rate of 1 × 10^−4^ and a momentum of 0.9. The stopping criterion was the number of epochs (15).

### Convolutional Deep Clustering Network (CDCN)

The convolutional neural network (KiwiNet) was jointly trained with a clustering/classification methodology (LAMDA) to generate a feature vector that accurately characterizes vocal individuality. The CNN is pre-trained to identify a set of individuals. Then, its latent space is used as the input feature vector to LAMDA, which clusters the data in online learning mode using the data labels to initialize the cluster prototypes. Next, clustering and CNN cross-entropy losses are combined and back-propagated throughout the CDCN using *L* = *αL_n_* + (1 – *α*)*L_c_* where *α* ∈ [0, 1] is a weighting factor, and *L_n_* and *L_c_* are the CNN and clustering losses, respectively. *α* = 0.5, i.e., equal weights for both losses was selected for this specific application. Once the training process has been completed, the training loops are disconnected and KiwiNet (trained) is used as the feature extractor for LAMDA, which performs clustering and classification tasks. The CDCN was trained using stochastic gradient descent with momentum and restarts. Momentum was chosen as 0.9. The learning rate (*LR*) was updated using: *LR*(*u*) = *LR*(0)/(1 + *d* * *p*). Where *d* = 0.01 is the learning rate decay, *LR*(0) = 1 × 10^−3^ is the initial learning rate, *u* is the current step, and *p* the mini-batch iteration number. LR was restarted every 5 epochs to *LR*(0). The training was performed in mini-batches of 16 calls, with calls being randomized every epoch, and the stopping criterion was the number of epochs (100).

### LAMDA (Learning Algorithm for Multivariate Data Analysis)

LAMDA (12, 13, 15, 16) is a fuzzy clustering and classification methodology. In addition to supervised learning, LAMDA allows: (i) online learning with unseen or novel class discovery (i.e., creation of classes that were not present during the training stage), and (ii) unsupervised learning without prior knowledge of the number of classes. These two features provide our framework with the ability to identify unexpected individuals and perform acoustic censusing.

LAMDA is not a distance-based method and uses adequacy degrees to estimate similarity among data. The contribution of each feature to a cluster is called the marginal adequacy degree (MAD). For each vocalization *v*, the MAD 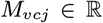 of each *j – th* feature to the cluster *c* is estimated using:

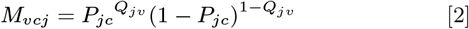

where 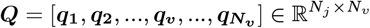, and 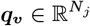 is a vector containing the normalized ([0, 1]) features of the call *v*. 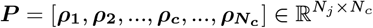 is a matrix with the cluster prototypes (means) of each feature *j* in each cluster *c*. *N_j_*, *N_c_*, and *N_v_* are the number of features, clusters, and vocalizations (calls), respectively.

Once MADs are estimated, these are combined using a fuzzy aggregation operator Eq. (3) in order to obtain a global adequacy degree (GAD, 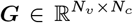), or membership degree, from an object (call) to each generated cluster (individual).

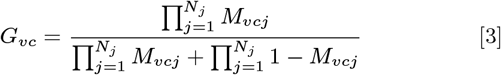

Since GADs are fuzzy elements ([0, 1]), hard (one-to-one) individual-call correspondences are determined by assigning a call to the individual with the highest membership degree.

The version of LAMDA used in this manuscript is called full reinforced LAMDA(16), and it uses a Yager aggregation operator Eq. (3) that naturally restricts the number of generated clusters (16, 31).

#### Individual Identification (Supervised learning)

In order to perform individual identification, LAMDA is used in supervised mode, where the number of individuals is known and cluster prototypes (***P***) are directly obtained by averaging the features using the data labels. Then call assignments are performed using Eqs. (2 and 3).

#### Unseen Individual Discovery (online learning)

Individuals not present in the training stage are discovered by allowing LAMDA to generate new clusters. To accomplish this, we add a Non-Informative Class (NIC), or class 0, to the cluster prototypes.

The algorithm operates as previously described using Eqs. (2 and 3), but when a call *v* is unrecognized (i.e., its maximum membership degree belongs to the NIC), a new cluster is created and initialized with the NIC parameters (*P*_*j*0_ = 0.5, ∀*j* = 1, …, *N_j_*) modified by the new data values (Eq. (4)). Note that 0.5 is a non-informative value for both MADs and GADs. if *P_jc_* = 0.5 then ∀ *Q_vj_* ∈ [0, 1], *M_vcj_* = 0.5 (Eq. (2)). Similarly for GADs, if *M_vcj_* = 0.5 ∀ *j* = 1, …, *N_j_*, then *G_vc_* = 0.5 (Eq. (3))

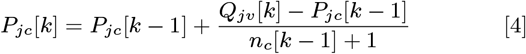

where 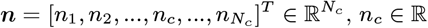 is the number of objects (calls) classified in the cluster *c*, and *k* is the current step.

In our specific application, we only allowed the algorithm to update the newly generated clusters, as the identification accuracy for the known individuals using the training data was highly accurate (> 90%). To make our framework insensitive to the order of cluster generation, we randomized the data before the clustering analysis and ran LAMDA 1000 iterations.

#### Acoustic Censusing (unsupervised learning)

For the censusing of individuals, LAMDA is used as a clustering algorithm (unsupervised mode). In this case, LAMDA starts with only one predefined cluster: the NIC, where the first call is always allocated. Then, a new cluster is created using Eq. (4), but its prototypes (*P*_*j*1_, ∀*j* = 1, …, *N_j_*) are initialized in the first step (*k* = 1) using the NIC parameters (*P*_*j*0_ = 0.5, ∀*j* = 1, …, *N_j_*) and *n_c_*[*k* – 1] = 1. After the first cluster has been created, the algorithm proceeds as disclosed in the online learning section, but allowing prototype modification for all clusters except the NIC.

### Cluster Validation

A cluster validation stage was implemented to automatically identify disjoint and sparsely-distributed clusters that could not be associated with specific individuals. To accomplish this, the quality of the clustering partition was evaluated using Eq. (5):

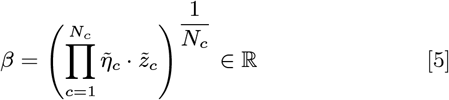

where 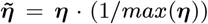 and 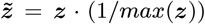. Here 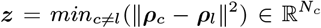 is a vector with the minimum squared distances between cluster prototypes, and 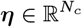 is a vector containing the number of objects in each cluster. *β* ∈ [0, 1] combines measures of cluster evenness 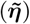 and separation 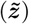, and is an estimator of cluster ’segregation’ in the partition. The lower the value of *β*, the more uneven and close-together the clusters are.

When the index *β* of the clustering partition is below its non-informative value (0.5), a criterion *τ* Eq. (6) automatically separates dense and compact clusters from disjoint and sparsely-distributed ones. *τ* is determined by Eq. (6):

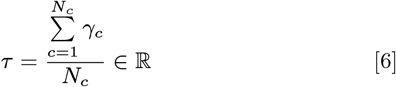

where, 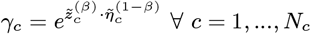 is the cluster’s dilated segregation. For any given cluster *c*, should *γ_c_* > *τ* the cluster is considered as an individual, otherwise it is categorized as a low-confidence cluster.

### Data Visualization

We used t-SNE (1500 iterations) for a visual representation of the calls in a two-dimensional space. Perplexity was selected as 35 for all figures. In Figs. 1C, 3A-B, and 4A-B the dimensionality reduction was performed on the feature set extracted via KiwiNet. t-SNE was used only for visualization and it was not part of the acoustic data analyses. We used the t-SNE Matlab implementation by van der Maaten and Hinton (33).

### Spectro-temporal features

Eight spectro-temporal call features were used for comparison with our proposed approach (Fig. 2A). Temporal features (i.e., call duration, syllable duration, intersyllable interval, number of syllables) were estimated finding the peaks, and their respective widths, of the averaged spectrogram in the spectral domain. The number of syllables is equivalent to the number of peaks, the syllable duration is the peak width at half height of peak prominence. The inter-syllable interval is the time period measured from the end of a syllable to the beginning of the next one, and the call duration the period from the beginning of the first syllable to the ending of the last one. The bandwidth is the spectrum between the frequencies at the half-power point, the dominant frequency is the frequency where most power is concentrated, and the spectral centroid is the frequency in which the centroid of the power spectral distribution is located, computed using 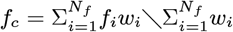, where 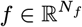 is a vector with the frequency values of each i-th bin of the spectrogram, and 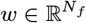 is the mean spectrum (i.e., the arithmetic mean of the spectrogram for all temporal values).

### Mel-frequency Cepstral Coefficients

MFCCs (Fig. 2B) were estimated in frames of 100 ms using a hamming window with frame shifts of 50 ms. In total, 40 coefficients, extracted from 36 filters, were used. Signals were pre-emphasized (0.99) before MFCC estimation and the coefficients lifted using a sinusoidal lifter of 25. The first MFCC contains the DC value and was not considered for the analysis. MFCCs were extracted using Wojcicki’s Matlab implementation (34).

### Software

All algorithms used in this manuscript were implemented in Matlab-R2021a.

## Author Contributions

C.L.B. and L.E.M. designed research. C.L.B. developed the computational and signal processing techniques and performed the data analysis. L.E.M. designed the ecological aspects of the experiment, and collected data. C.L.B and L.E.M. wrote the manuscript.

## ACKNOWLEDGMENTS

We thank Jennifer Dent, Vanessa Mander, Peter Jahn, Ray Beckford, Kristy Owens, Jo Halley, Sandy Yong, and Kaitlin Morrison for their fieldwork (recorder deployment/maintenance and location of nests). We also thank the NZ Department of Conservation (NZDOC), the Paparoa Wildlife Trust (PWT), Ngāi Tahu, and Roa Mining Company for sharing, and/or facilitating the collection of, acoustic data. We are grateful to NZDOC, Lincoln University’s Department of Pest Management and Conservation, and PWT for loan of acoustic recorders. This project was financially supported by Verum Group (a sister company to Atarau Sanctuary).

## Appendix

**Figure S1:**
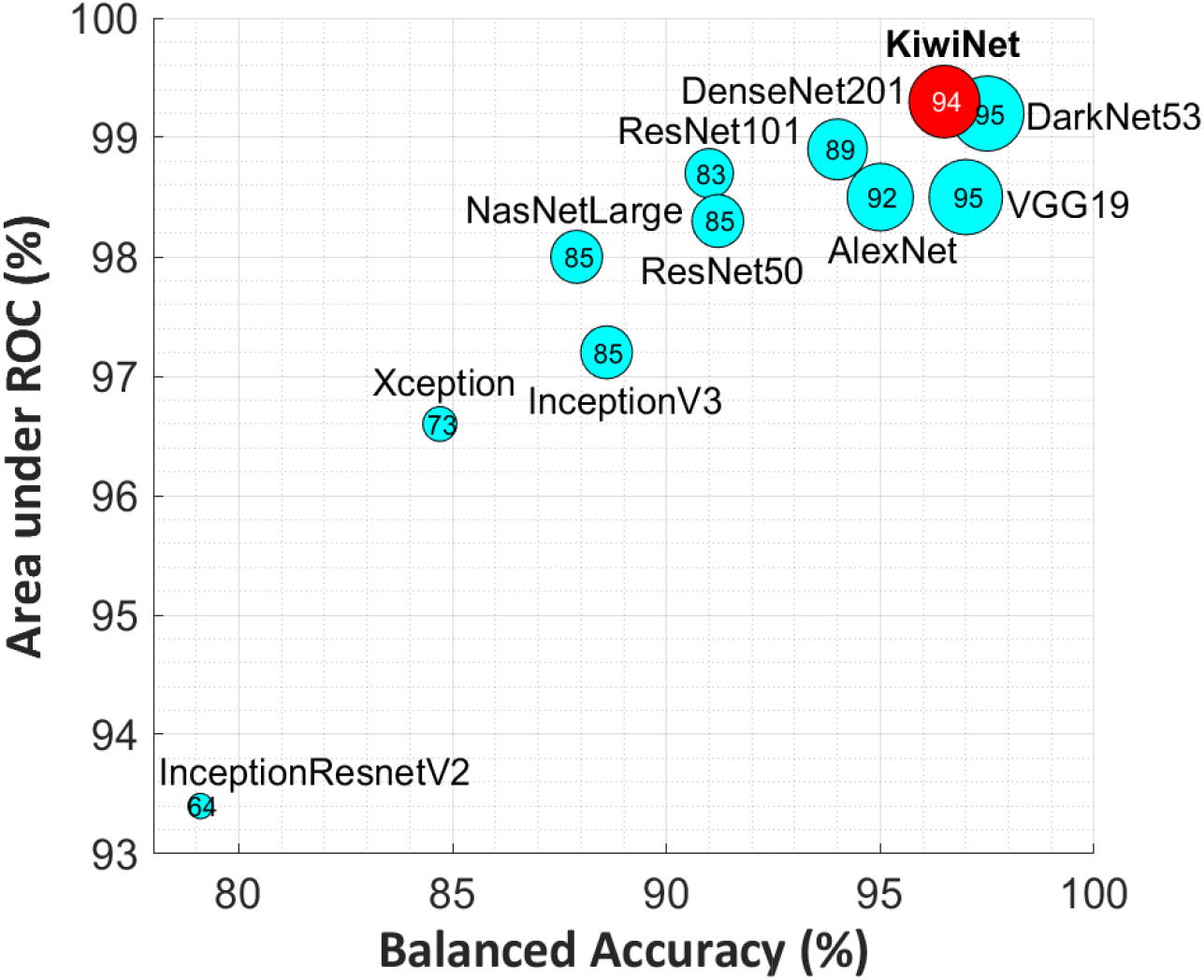
Comparison of KiwiNet with commonly-used CNN Architectures. Comparison of our proposed convolutional Neural Network (CNN) with other frequently-used CNN architectures. All Networks were trained to recognize 10 individuals (5♂and 5♀). The size of the markers and the number inside them represent the *F*_1_-score = 2 (precision × sensitivity) / (precision + sensitivity). y-axis is the average area under the receiver operating characteristic (ROC) curve and the x-axis is the average balanced accuracy ((sensitivity + specificity) / 2) for all individuals. All networks were pre-trained using the imagenet dataset.

**Figure S2:**
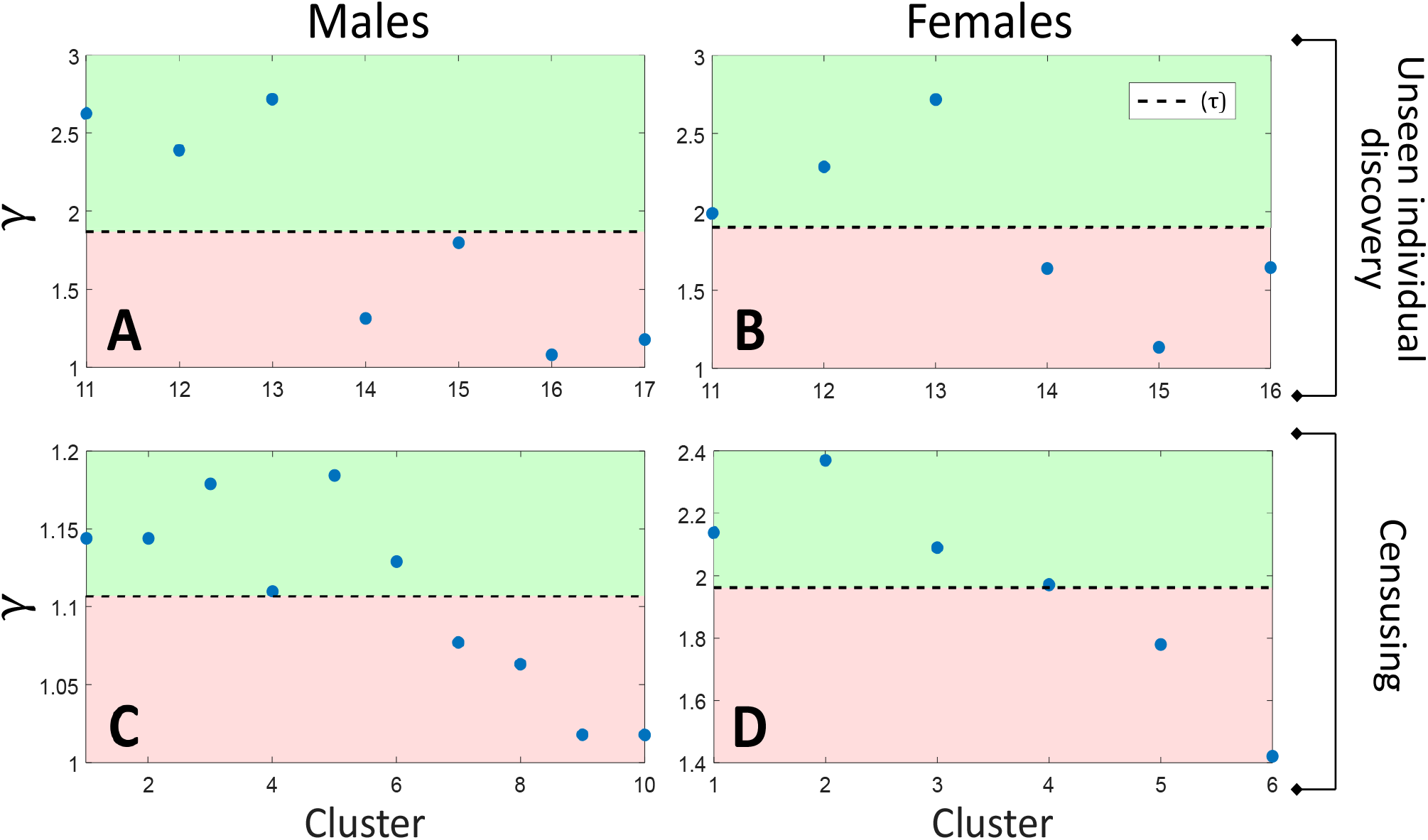
Cluster Validation. Cluster validation for the unseen individual discovery (**A**,**B**; see Fig. 3 of the manuscript) and acoustic censusing (**C**,**D**; see Fig. 4 of the manuscript) tasks. Dotted lines are the means (*τ*) of ***γ*** (Eq. 6 in the manuscript). Green region (acceptance) indicates clusters categorized as individuals. Red region (rejection) indicates low-confidence clusters. (**A**,**C**) and (**B**,**D**) represent data for males and females, respectively.

**Table S1:** Detailed architecture of KiwiNet.

|  | Layer Type | Activations | Parameters |
| --- | --- | --- | --- |
| 1 | Image Input | 224x224x3 | Zero-center normalization |
| 2 | Convolution | 224x224x64 | 64 3x3x3 convolutions with stride [1 1] and padding [1 1 1 1] |
| 3 | ReLU | 224x224x64 |  |
| 4 | Convolution | 224x224x64 | 64 3x3x64 convolutions with stride [1 1] and padding [1 1 1 1] |
| 5 | ReLU | 224x224x64 |  |
| 6 | Max Pooling | 112x112x64 | 2x2 max pooling with stride [2 2] and padding [0 0 0 0] |
| 7 | Convolution | 112x112x128 | 128 3x3x64 convolutions with stride [1 1] and padding [1 1 1 1] |
| 8 | ReLU | 112x112x128 |  |
| 9 | Convolution | 112x112x128 | 128 3x3x128 convolutions with stride [1 1] and padding [1 1 1 1] |
| 10 | ReLU | 112x112x128 |  |
| 11 | Max Pooling | 56x56x128 | 2x2 max pooling with stride [2 2] and padding [0 0 0 0] |
| 12 | Convolution | 56x56x256 | 256 3x3x128 convolutions with stride [1 1] and padding [1 1 1 1] |
| 13 | ReLU | 56x56x256 |  |
| 14 | Convolution | 56x56x256 | 256 3x3x256 convolutions with stride [1 1] and padding [1 1 1 1] |
| 15 | ReLU | 56x56x256 |  |
| 16 | Convolution | 56x56x256 | 256 3x3x256 convolutions with stride [1 1] and padding [1 1 1 1] |
| 17 | ReLU | 56x56x256 |  |
| 18 | Convolution | 56x56x256 | 256 3x3x256 convolutions with stride [1 1] and padding [1 1 1 1] |
| 19 | ReLU | 56x56x256 |  |
| 20 | Max Pooling | 28x28x256 | 2x2 max pooling with stride [2 2] and padding [0 0 0 0] |
| 21 | Convolution | 28x28x512 | 512 3x3x256 convolutions with stride [1 1] and padding [1 1 1 1] |
| 22 | ReLU | 28x28x512 |  |
| 23 | Convolution | 28x28x512 | 512 3x3x512 convolutions with stride [1 1] and padding [1 1 1 1] |
| 24 | ReLU | 28x28x512 |  |
| 25 | Convolution | 28x28x512 | 512 3x3x512 convolutions with stride [1 1] and padding [1 1 1 1] |
| 26 | ReLU | 28x28x512 |  |
| 27 | Convolution | 28x28x512 | 512 3x3x512 convolutions with stride [1 1] and padding [1 1 1 1] |
| 28 | ReLU | 28x28x512 |  |
| 29 | Max Pooling | 14x14x512 | 2x2 max pooling with stride [2 2] and padding [0 0 0 0] |
| 30 | Convolution | 14x14x512 | 512 3x3x512 convolutions with stride [1 1] and padding [1 1 1 1] |
| 31 | ReLU | 14x14x512 |  |
| 32 | Convolution | 14x14x512 | 512 3x3x512 convolutions with stride [1 1] and padding [1 1 1 1] |
| 33 | ReLU | 14x14x512 |  |
| 34 | Convolution | 14x14x512 | 512 3x3x512 convolutions with stride [1 1] and padding [1 1 1 1] |
| 35 | ReLU | 14x14x512 |  |
| 36 | Convolution | 14x14x512 | 512 3x3x512 convolutions with stride [1 1] and padding [1 1 1 1] |
| 37 | ReLU | 14x14x512 |  |
| 38 | Max Pooling | 7x7x512 | 2x2 max pooling with stride [2 2] and padding [0 0 0 0] |
| 39 | Convolution | 4x4x32 | 32 1x1x512 convolutions with stride [2 2] and padding [0 0 0 0] |
| 40 | Global Average Pooling | 1x1x32 |  |
| 41 | Fully Connected | 4096 | 4096 |
| 42 | ReLU | 4096 |  |
| 43 | Dropout | 4096 | 50% |
| 44 | Fully Connected | 4096 | 4096 |
| 45 | ReLU | 4096 |  |
| 46 | Dropout | 4096 | 50% |
| 47 | Fully Connected | # of Individuals |  |
| 48 | Softmax | # of Individuals |  |

**Table S2:** Classification results for individual identification and unseen individual discovery.

| **Males** | Acc* (%) | F1** (%) | **Females** | Acc (%) | F1 (%) |
| --- | --- | --- | --- | --- | --- |
| Male 1 | 87.5 | 85.7 | Female 1 | 100.0 | 100.0 |
| Male 2 | 98.8 | 70.6 | Female 2 | 100.0 | 100.0 |
| Male 3 | 99.8 | 85.7 | Female 3 | 99.7 | 85.7 |
| Male 4 | 65.9 | 36.4 | Female 4 | 100.0 | 100.0 |
| Male 5 | 99.5 | 80.0 | Female 5 | 87.5 | 85.7 |
| Male 6 | 100.0 | 100.0 | Female 6 | 89.7 | 80.0 |
| Male 7 | 79.0 | 60.0 | Female 7 | 100.0 | 100.0 |
| Male 8 | 80.8 | 63.6 | Female 8 | 98.9 | 50.0 |
| Male 9 | 92.1 | 89.2 | Female 9 | 82.5 | 50.0 |
| Male 10 | 90.4 | 86.7 | Female 10 | 87.5 | 85.7 |
| Unknown male 1 | 87.5 | 81.8 | Unknown female 1 | 80.4 | 74.2 |
| Unknown male 2 | 79.3 | 73.9 | Unknown female 2 | 83.2 | 67.7 |
| Unknown male 3 | 88.3 | 84.0 | Unknown female 3 | 76.3 | 69.0 |

\*Balanced Accuracy = (sensitivity + specificity) / 2, \*\* $F_1$ -score = 2 (precision × sensitivity) / (precision + sensitivity).

**Table S3:** Censusing Accuracy.

| Males | Acc* (%) | F1** (%) | Females | Acc (%) | F1 (%) |
| --- | --- | --- | --- | --- | --- |
| Male 1 | 92.6 | 90.9 | Female 1 | 85.9 | 80.0 |
| Male 2 | 83.7 | 80.5 | Female 2 | 93.5 | 80.0 |
| Male 3 | 93.3 | 88.0 | Female 3 | 75.1 | 67.3 |
| Male 4 | 81.7 | 76.7 | Female 4 | 100.0 | 100.0 |
| Male 5 | 97.2 | 96.2 |  |  |  |
| Male 6 | 80.7 | 71.9 |  |  |  |
| Average | 88.2 | 84.0 |  | 88.6 | 81.8 |

\*Balanced Accuracy = (sensitivity + specificity) / 2, \*\*F<sub>1</sub>-score = 2 (precision × sensitivity) / (precision + sensitivity).

